# Exhaustive search for DNA binding sites of *Basillus subtilis* transcription factor YsiA by KaScape method

**DOI:** 10.1101/2023.10.16.562645

**Authors:** Bing-yao Huang, Mia Huang, Hong Chen, Xiao-dong Su

**Affiliations:** State Key Laboratory of Protein and Plant Gene Research, School of Life Sciences, and Biomedical Pioneering Innovation Center (BIOPIC), Peking University, Beijing, 100871, China; Grasse Town No. 2015, Tongzhou District, Beijing 101119, China

## Abstract

*Basillus subtilis* (*B. subtilis*) YsiA is also known as FadR is a pre-formed dimeric repressor to regulate fatty acid degradation. However, the knowledge about double-stranded DNA(dsDNA) binding sites of YsiA in the whole genome level is limited so far. In this report, we have applied a newly established high-throughput method, KaScape, to exhaustively study the relative binding sites and energy landscape of YsiA and confirmed the resuls with gel-shift experiments. We’ve found that half-binding site is enough for YsiA to bind with dsDNA. Besides detecting the known consensus sequence which is bound by YsiA, we have also detected other minor binding site sequences through the KaScape method. It thus demonstrates that KaScape can be used to study dimeric transcription factors (TFs) in general.

## 1. Introduction

In bacteria, fatty acids are essential components in their metabolism, cell membrane, metabolic energy, and overall homeostasis. The amount of fatty acids is regulated through production and degradation paths. In general, it possesses two global transcription regulators in most bacteria, FadR (also known as YsiA *B. subtilis*) and FapR. Briefly, YsiA mainly regulates the fatty acid degradation, whereas FapR regulates fatty acid production (1). We would focus on the double strand DNA (dsDNA) binding behavior of YsiA in this work.

Through the framework of the Japan Functional Analysis Network for *B. subtilis* and DNA microarray analyses, researchers have found that YsiA protein negatively regullates more than 10 genes, most of which participate in fatty acid β-oxidation process (a key process to fatty acid degradation) which leads to the discovery of YsiA’s central regulatory role in fatty acid degradation (2). The YsiA protein is a member of the TetR family with a bacterial helix-turn-helix fold (3) . It is a pre-formed dimeric repressor and can bind to a consensus pseudo-palindromic sequence WTGAATGAMTANTCATTCAN (2). Through the FadR-dsDNA complex crystal structures, we can see that the YsiA half-binding site which is defined as the binding site of one monomer in the dimeric form is ATGAAT (4).

The current binding sequences information of YsiA is mainly from the microarray data (2). Howerver, the sequence amount tested is limited, and the microarray data has the potential to present false positives or results with low accuracy (5). In order to extensively study the relative binding affinity of YsiA, we have used the KaScape method (6) as our main form of analysis to study the dsDNA binding characteristic of YsiA. This approach enables us to inspect all kinds of dsDNA to protein binding, under identical conditions. The KaScape method can successfully lead us to capture diverse binding interactions in the dsDNA pool as well as the protein YsiA interaction. It can provide us with a higher probability of encountering unique DNA sequences that exhibit a different binding characteristic from what we already understand with the protein YsiA.

We have undertaken a series of KaScape experiments with random base lengths ranging from 4 to 7. After data analysis, we found that the results of the experiments with random base lengths of 4, 5, 6, and 7 are highly consistent. We also confirmed the KaScape results by running several sequences through electrophoretic mobility shift assay (EMSA) analysis. The KaScape results can also predict the binding capability of the known consensus sequence for YsiA. We found the half-binding site is enough for YsiA to bind and we’ve found other bound sequences for YsiA besides the consensus sequences.

## 2. Materials and methods

### 2.1 Random dsDNA preparation

Random dsDNA pool was prepared according to the procedures detailed in the research by Hong Chen et al (6).

### 2.2 Protein preparation

The *B. subtilis* YsiA protein used in this study was prepared using *Escherichia coli* BL21 as previously reported (7). In brief, the YsiA gene from *B. subtilis* was subcloned by polymerase chain reaction (PCR) for expression in *E. coli* and inserted into the pET21b vector along with a C-terminal 6xHis-tag. The transformed *E. coli* colonies were used to inoculate LB culture supplemented with 0.4% glucose, grown overnight at 37°C. The cells were then used to inoculate 2-L cultures and induced by adding isopropyl beta-D-1-thiogalacto-pyranoside (IPTG) to a final concentration of 0.4 mM when they reached an OD_600nm_ of 0.6-0.8. The cells were then grown for an additional 18 hours at 16°C and harvested. The centrifuged and collected cells were resuspended in a buffer of 20 mM Tris (pH 8.0) and 100 mM NaCl (30 mL per Liter of cells). They were then stored at -80°C for further purification.

To purify the YsiA protein, cells were sonicated and centrifuged, and then they were resuspended in buffer of 20 mM Tris (pH 8.0) and 100 mM NaCl. The cells were then centrifuged at 32,000g at 4°C and the supernatant was collected.

The supernatant containing the target protein was then loaded on a 5 mL Ni-NTA column followed by a size exclusion column (Superdex 75, GE Healthcare, USA). YsiA is then eluted stepwise with 30 mM, 60 mM, 90 mM, 120 mM, 150 mM, 210 mM, and 300 mM imidazole after the supernatant is washed with a buffer containing 20 mM Tris (pH 8.0), 100 M NaCl, 10 mM imidazole, and 2 mM β-mercaptoethanol. The purity of the YsiA proteins was confirmed by sodium dodecyl sulfate polyacrylamide gel electropheresis (SDS-PAGE) and then concentrated to around 2 mg/mL using an MWCO 10kDa centrifugal concentrator.

### 2.3 KaScape procedures

The exhaustive search of the binding sequence for YsiA was conducted using KaScape method, following the steps detailed in Hong Chen, et al.(6). Briefly, a random dsDNA pool containing 4 bp, 5 bp, 6 bp, or 7 bp was prepared and mixed with purified His-tagged YsiA for 30 minutes. Then the protein-DNA binding complex was separated using His-tag magnetic beads. The bound dsDNA was then purified using the Oligo Clean & Concentrator Kits (ZYMO Research, USA). To get the bound dsDNA ready for sequencing, they were extended to 75 bp by PCR. The original random dsDNA pool was also extended to 75 bp by PCR. The extended dsDNA oligos were then purified using DNA Clean & Concentration Kits (ZYMO Research, USA). The purified and extended dsDNA were then ligated to Illumina sequencing adaptors and extended, then further sequenced by Illumina NovaSeq PE150 platform. The sequencing data was analyzed and the K-mer graph was drawn.

### 2.4 KaScape data analysis

The relative binding energy calculation method is a little bit different from the original KaScape paper (6). For each random base length (n) KaScape experiment, the relative binding energy landscape with a series of short sequence length (m) can be calculated. The short sequence length ranges from 3 bp to n bp. The number of reads for each short sequence type, *s*_*i*_, in the random dsDNA pool sequencing library was counted as 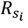. For the remaining reads of the sequencing results obtained from the bound dsDNAs, the number of reads for each short sequence type, *s*_*i*_, in the bound dsDNAs sequencing library was counted as 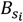. Then, the proportion of the short sequence type *s*_*i*_ in the random dsDNA pool was calculated as 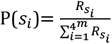 ; while the proportion of the short sequence type *s* in the bound dsDNAs was calculated as 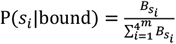. Finally, the relative binding energy which represents the affinity (8) of short sequence *s*_*i*_ was calculated as 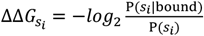. The above data analysis was performed using custom code written in Python 3.6. The following modules were used: re, os, sys, datetime, collections, gc, pandas, numpy, math, matplotlib, pickle, and xlwt.

### 2.5 Electrophoretic mobility shift assay (EMSA)

The reverse complementary ssDNAs were annealed to form dsDNA. The typical binding reaction (6 ul) contained dsDNA (6 pmol), 25 mM HEPES (pH 7.0), 100 mM NaCl, 5% glycerol, and proteins. The binding reaction mixture was incubated at 4°C for 20 minutes, and the complex was loaded on a 10% nondenaturing polyacrylamide gel electrophoresis (PAGE) in 0.5×TBE at 120 V for 1 hour. Lastly, the DNA was dyed and detected using AI600 (GE Healthcare, USA).

## 3. Results and discussions

### 3.1 The purification of YsiA

In order to extensively explore the relative binding energy landscape for YsiA, gaining the highly purified YsiA is the essential first step. The preparation method of YsiA is mentioned in the Protein preparation section (see 2.2). Fig. 1 shows the purification results of the Ni-chelating column and size-exclusive chromatography. The YsiA protein is eluted at 160 mM imidazole (32%). The molecular weight of the YsiA DBD is 23 kDa. The peak volume in the size-exclusive chromatography map is 13.95 mL which suggests the free YsiA forms dimer in solution. The SDS-PAGE results showed that the YsiA is highly purified with a large amount (around 50 mg).

**Fig. 1.**
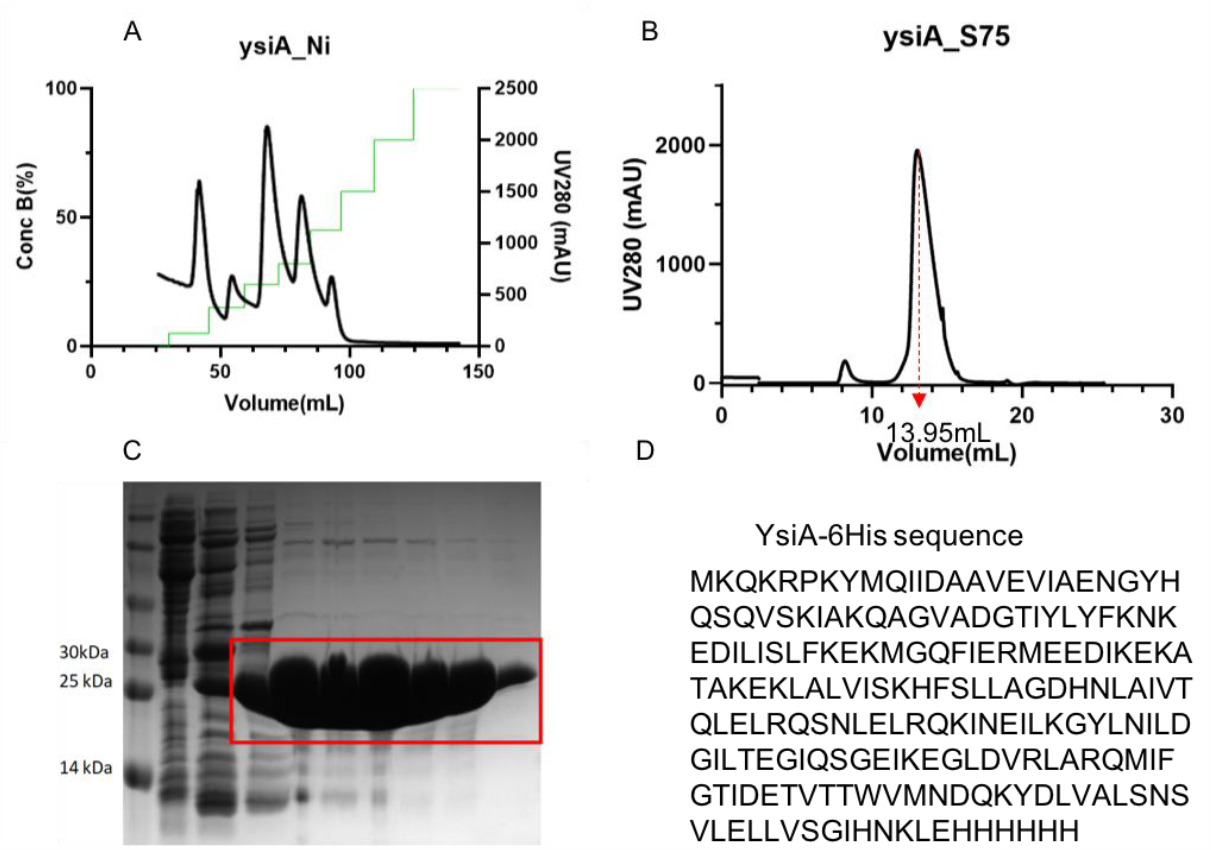
The purification of the YsiA protein. (A) Ni-chelating column purification result. (B) size-exclusive chromatography result. (C) SDS-PAGE result for Ni-chelating column purification samples. (D) the YsiA DBD sequence.

### 3.2 Binding affinity landscape for YsiA

We carried out a series of KaScape experiments for YsiA. The random base length of the random sequence used in the KaScape experiment is 4, 5, 6, or 7 bp, respectively. After sequencing data analysis, we calculated the relative binding energy landscape for short sequences ranging from 3 bp to n bp, see Fig. 2. The relative binding landscape maps from each row are calculated from one random base length type of KaScape experiment, respectively. Most of the pearson correlation coefficient between the KaScape experiment is higher than 0.5 which means the KaScape experiment results are consistent and robust (see Fig. 2 and Fig. 3). In the introduction part, we have mentioned that YsiA can bind with a consensus sequence WTGAATGAMTANTCATTCAN, where W, M, and N stand for A or T, A or C, and any base, respectively (2). The consensus sequence is reverse complementary from each other which is a typical characteristic of dimer transcription factors. By analyzing the YsiA and dsDNA complex structure (4), we’ve found the half site bound by YsiA (that is the YsiA monomer binding site) is ATGAAT. From the KaScape experiment where the random length equals 6, the relative binding energy of ATGAAT is -0.3702 whereas the relative binding energy of TTGAAT is - 0.2830. The relative binding energy predicted by KaScape experiment is less than 0 which means those sequences are the binding signal.

**Fig. 2.**
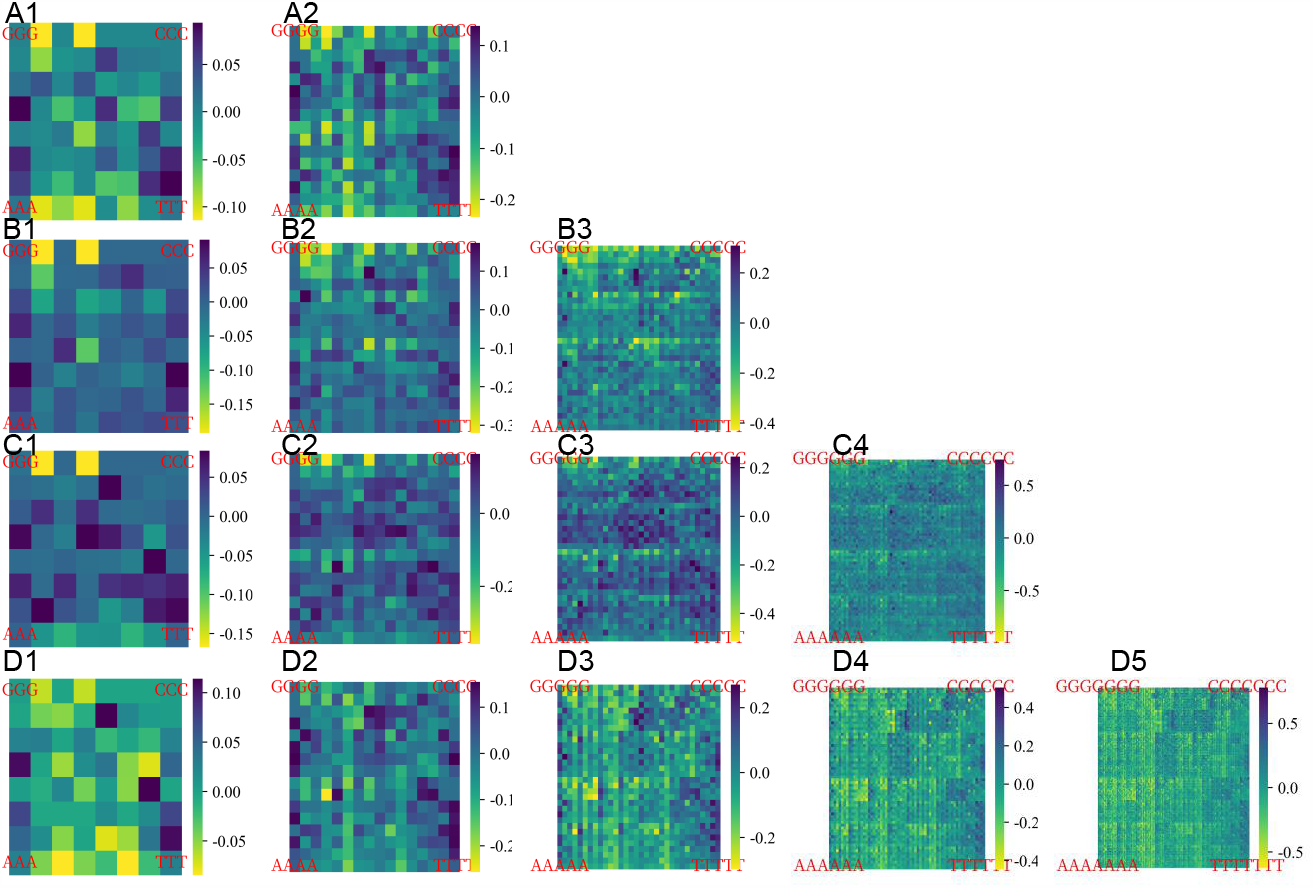
The relative binding energy landscape for YsiA for each short sequence length. Each row represents one KaScape experiment with a certain random base length. For the first row, the random base length of the random sequence used in the KaScape experiment is 4. For the second row, the random base length of the random sequence used in the KaScape experiment is 5. For the third row, the random base length of the random sequence used in the KaScape experiment is 6. For the fourth row, the random base length of the random sequence used in the KaScape experiment is 7. Each column represents one short sequence length type. For the first column, the short sequence length is 3. For the second column, the short sequence length is 4. For the third column, the short sequence length is 5. For the fourth column, the short sequence length is 6. For the fifth column, the short sequence length is 7.

**Fig. 3.**
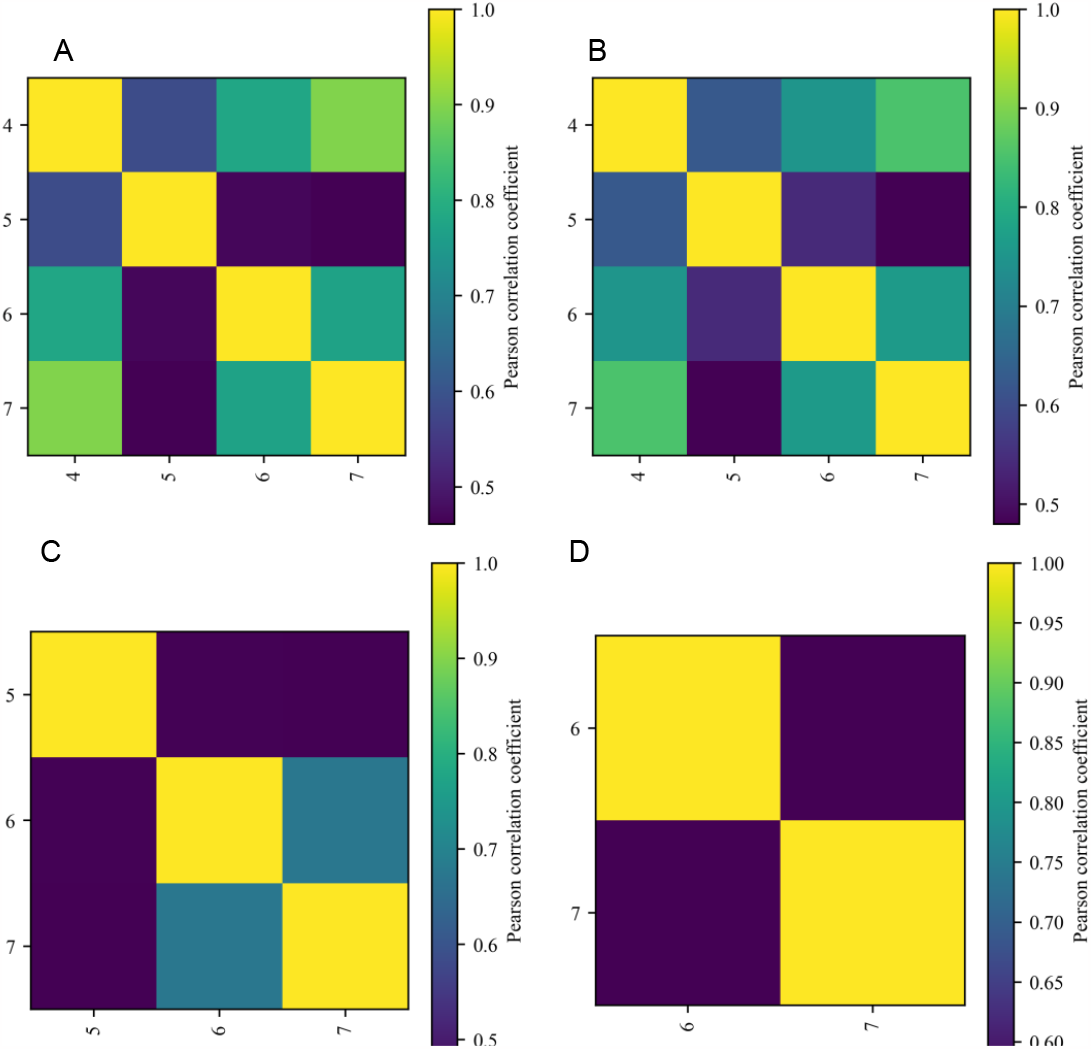
The consistency of the KaScape experiment for different short sequence length maps. The x- and y-axis labels represent the random base length used in the KaScape experiment. (A) the short sequence length of the relative binding energy maps calculated for the pearson correlation coefficient is 3. The relative binding energy maps are those in the first column in Fig. 2. (B) the short sequence length of the relative binding energy maps calculated for the pearson correlation coefficient is 4. The relative binding energy maps are those in the second column in Fig. 2. (C) the short sequence length of the relative binding energy maps calculated for the pearson correlation coefficient is 5. The relative binding energy maps are those in the third column in Fig. 2. (D) the short sequence length of the relative binding energy maps calculated for the pearson correlation coefficient is 6. The relative binding energy maps are those in the fourth column in Fig. 2.

### 3.3 The binding behavior of YsiA revealed by KaScape and EMSA

To see whether the relative binding energy predited by the KaScape experiment is correct and to extensively explore the sequences bound by YsiA besides the known consensus sequence, we did the EMSA for several sequences and compare the results with the relative binding energy predicted by KaScape experiment (see Fig. 4). We found the KaScape prediction results are consistent with EMSA result. Besides the consensus sequence which can be bound by YsiA (see the sequence 1 in Fig. 4A), The YsiA can bind other sequences. The half-site is enough for YsiA to bind dsDNA (see sequence 2, 5 and 6 in Fig. 4A which only contain one ATGAAT). In the half-site ATGAAT, TGA is enough for YsiA to bind dsDNA (compare the sequences 3, 4, 5, and 7 in Fig. 4A). Besides the consensus half-site, the YsiA can bind other sequences (see sequences 7, 8, and 9 in Fig. 4A).

**Fig. 4.**
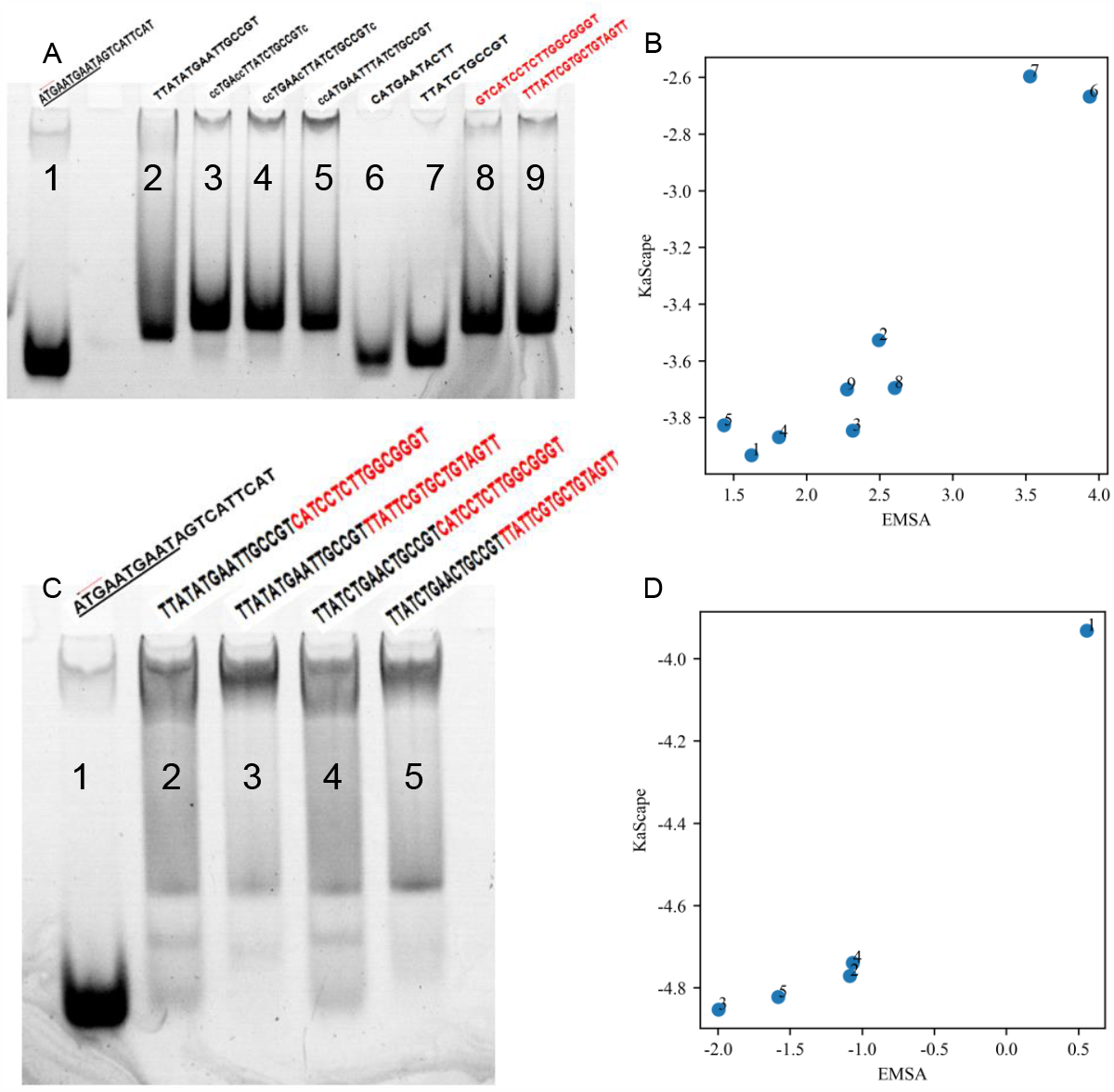
The EMSA results and the consistency between KaScape results and EMSA. The molar ratios of YsiA to DNA is 4:1. (A) and (C) are the EMSA result for several sequences which is annotated in the above line. (B) and (D) the relative binding energy comparison between the KaScape experiment and EMSA. The KaScape value is the relative energy value in each sequence which add up all the 6-mer association constant Ka of the sequence from the KaScape result, and then calculate the relative binding energy as -log2(Ka) The EMSA value is calculated as -log2(bound/unbound). The bound and unbound are the mean grayscale values in the bound and unbound region quantified from the EMSA experiment. (B) The sequences are listed in (A). The number order is the same as the sequence order in (A). (D) The sequences are listed in (C). The number order is the same as the sequence order in (C).

The association constant of the long sequence can be added up by the association constant of the short sequence. The red part of the sequences 2 and 4 in Fig. 4 C are the same as the sequence 6 in Fig. 4 A. The red part of the sequences 3 and 5 in Fig. 4 C are the same as the sequence 7 in Fig. 4 A. The black part of sequences 2 and 3 are the same which contain the consensus half-site ATGAAT whereas the black part of sequences 4 and 5 are the same which contain the CTGAAC. The relative binding energy of sequence 7 is lower than sequence 6 shown in Fig. 4 A and Fig. 4 B. That means the relative binding energy of the red part of sequences 3 and 5 is lower than sequences 2 and 4 in Fig. 4 C. The relative binding energy of ATGAAT (-0.2547, predicted by KaScape) is lower than CTGAAC (0.0603, predicted by KaScape). This means the relative binding energy of the black part of sequence 3 (or 2) is lower than sequence 5 (or 4) in Fig. 4 C. The difference in the relative binding energy of the black part is larger than the red part, so the relative binding energy order is 3 < 5 < 2 < 4 in Fig. 4 D.

## Conclusions

We have taken YsiA as an example to demonstrate that KaScape experiments can be used to study pre-formed dimeric transcription factors. We have done a series of KaScape experiments with random base lengths ranging from 4 to 7 for *B. subtilis* Ysia. The series of KaScape results are consistent with each other and the results is confirmed by EMSA which shows the robustness and repeatability of the KaScape experiments.

From the KaScape experiment, we can not only obtain the consensus binding sequences but also other sequences bound by YsiA. Through searching the *B. subtilis’s* genome, and calculate the relative binding energy using the KaScape data, we may predict some more genes besides those used in fatty acid degradation that can be regulated by YsiA, and it can help us to broaden the functional roles of YsiA, thus to form an enriched gene regulation network. We also found that the half-binding site is enough for YsiA to bind which may be useful in understanding the searching and binding behavior of the pre-dimeric transcription factors.

## Author contributions

X.-D.S. conceived and supervised the entire project. B. H. mainly performed the experiments, M. H. participated and helped with the experiments, H.C. performed all the computational analyses and programming, X.-D.S., H.C., and M. H. discussed the data and wrote the manuscript.

